# The metabolic costs of meiotic drive

**DOI:** 10.1101/2024.10.24.620073

**Authors:** Sasha L Bradshaw, Enrique Rodriguez, Hanting Wang, Cyn Thea Yu, Cyril De Villiers De La Noue, Aadil Hafezjee, Andrew Pomiankowski, M. Florencia Camus

## Abstract

Selfish genetic elements, such as meiotic drive genes, disrupt Mendel’s law of equal segregation by biasing their own transmission, often at a detriment to the rest of the genome. The Malaysian stalk-eyed fly (*Teleopsis dalmanni)* sex-ratio (SR) meiotic drive system is located within a series of large inversions on the X chromosome subject to low recombination and is associated with deleterious effects on fitness. Here we examine the metabolic effects of meiotic drive across male and female stalk-eyed flies. High-resolution O2k respirometry coupled with whole-organism respirometry were used to obtain mitochondrial function and metabolic rates. Complimentary assays on food consumption established downstream effects of metabolism on nutrient acquisition. The experiments demonstrate that individuals with SR meiotic drive elements have impaired mitochondrial function and reduced capacity for ATP synthesis, as shown by a lower respiratory control ratio and weaker contribution of Complex I to respiration. Drive individuals also exhibited an increased basal metabolic rate and consumed a greater amount of food than wild-type individuals. These findings show that the drive genotype imposes metabolic costs in both male and female hosts. The disruption in mitochondrial function likely leads to compensation via an increase in both basal metabolic rate and nutrient acquisition. A potential cause lies in the accumulation of deleterious mutations in the inversions on the X chromosome that house the meiotic drive, which are subject to weak natural selection. In females, the drive chromosome has a dominant effect, with a single copy causing substantial metabolic compromise. There was little evidence of male-specific metabolic costs, nor evidence of an accumulation of sexually antagonistic effects of drive chromosomes on female metabolism. These results suggest that direct metabolic costs from meiotic drive on spermatogenesis and from sexually antagonistic selection are relatively weak. This research provides new insight into the interplay between meiotic drive and metabolism, drawing attention to the broader physiological repercussions selfish genetic elements may have on their hosts.

## 1. Introduction

Meiotic drivers are one of the most well-studied classes of selfish genetic elements that break Mendel’s law of equal segregation (Lindholm et al., 2016). They have been identified on the autosomes and the sex chromosomes—the latter being well-documented owing to the resulting bias in progeny sex ratio (Hurst et al., 1991; Jaenike, 2001; Sutter et al., 2015). Meiotic drive occurs via the preferential segregation toward the egg nucleus away from the polar bodies in females or through disabling non-carrier gametes post-meiotically in males to facilitate a transmission advantage of the drive allele (Courret et al., 2019). This loss of sperm typically results in direct costs to the organism because of the dysfunction or loss of gametes (Price et al., 2008; Sutter et al., 2015; Zanders et al., 2019). Fertility reduction is particularly strong under conditions of sperm competition, both in defensive and offensive roles (Atlan et al., 2004; Price et al., 2008).

A range of indirect organismal costs are associated with male meiotic drive (Lindholm et al., 2016). Loss of viability has been demonstrated in a range of species, both in females homozygous and heterozygous for drive alleles, as well as in drive-carrying males (Curtsinger et al., 1980; Dyer et al., 2007; Finnegan et al., 2019; Larner et al., 2019). These costs are thought to originate from the notion that drivers tend to position within low-frequency chromosomal inversions or other genomic regions subject to reduced recombination (Jaenike, 2001), resulting in weak selection and the accumulation of a high deleterious mutational load (Berdan et al., 2023; Kirkpatrick, 2010). In addition, the hemizygosity of the X chromosome means that recessive effects are selected more strongly in males than in females (both for and against) until homozygotes become more common (Carazo, 2022; Gibson et al., 2002; Innocenti et al., 2010; Ruzicka et al., 2019). This is predicted to lead to the enrichment of sexually antagonistic alleles on driving X chromosomes that benefit male over female fitness, with the greatest loss to female fitness in homozygotes (Rydzewski et al., 2016).

The magnitude and direction of direct and indirect costs of meiotic drive and their associated inversions have rarely been subjected to explicit empirical investigation. A means to gauge these costs is to examine the effects on metabolism (David et al., 2004; Shaar-Moshe et al., 2019). Metabolism and metabolic traits are fundamental to life history trait evolution, capturing the rate at which organisms transform and expend finite energy into growth, maintenance and reproduction (Jonsson et al., 2016; Larivée et al., 2010; Sinclair, 2015). Furthermore, metabolism is a useful “intermediate phenotype”, positioned between the level of the genotype and traditionally measured morphological and behavioural traits that contribute to fitness (Cameron et al., 2024; Pettersen et al., 2018). Although the association of metabolic traits with fitness is inevitably complex, it can provide a clear indication of where physiological processes are compromised by genetic variants and are, therefore, likely to reduce fitness (Blackmer et al., 2005; Criscuolo et al., 2008; Konarzewski et al., 2013; Mueller et al., 2001; Pettersen et al., 2016). An advantage to studying metabolic traits is that they can be assessed in an equivalent manner in both sexes, allowing the investigation of sexual differences, which is key to the hypotheses investigated here.

In this study, we measure three metabolic-related traits to evaluate how meiotic drive directly in males and indirectly through its genomic architecture (through associated chromosomal inversions) in both sexes leads to costs linked to downstream consequences for fitness. These are investigated in the Malaysian stalk-eyed fly, *Teleopsis dalmanni*, which carries an X-linked meiotic drive system (Bradshaw et al., 2022; Cotton et al., 2014; Cotton et al., 2010; Finnegan, et al., 2019; Meade et al., 2020; Meade et al., 2019). This species is known for its extreme sexual dimorphism, in which males have greatly exaggerated eyespans that are subjected to strong female mate preferences (Burkhardt et al., 1988; Wilkinson et al., 1998). Males that carry the meiotic drive X chromosome, known as *Sex-Ratio* or SR, cause the dysfunction of Y-bearing sperm and sire female-only broods (Atlan et al., 2004; Beckenbach, 1983; Beckenbach, 1978; Courret et al., 2019; Larracuente et al., 2012). SR meiotic drive appears to be broadly stable across natural populations at a frequency of around 20% (Paczolt et al., 2017; Presgraves et al., 1997). It is also present in the sister species *T. whitei,* which suggests a common origin estimated at around 4 million years ago (Paczolt et al., 2017; Wilkinson et al., 2003).

*T. dalmanni* drive males curiously do not suffer from reduced fertility despite the destruction of half of their gametes (Meade et al., 2020). They pass on the same number of sperm per ejaculate as wild-type males and have equal fertility when in competition with wild-type males (Bates et al., 2023; Meade et al., 2019). This can be explained by the adaptive enlargement of testes in drive males, which is evident in the primordia of the adult fly testes upon eclosion from the pupal stage, the accelerated growth rates during early adult development, and the increased size of testes in sexually mature flies (Bradshaw et al., 2022). A number of other traits associated with the meiotic drive X chromosome are thought to be detrimental to fitness. Drive males have reduced eyespan (Finnegan et al., 2021; Finnegan et al., 2019; Meade et al., 2020), smaller accessory glands (Meade et al., 2020)—though this is not well established, see (Bradshaw et al., 2022)—and mate less frequently than wild-type males (Finnegan et al., 2019). Drive females do not have reduced eyespan (Cotton et al., 2014; Finnegan et al., 2021) but have lower fecundity compared to wild-type females (Bates, 2023). Both sexes have reduced egg-to-adult viability when they carry the drive chromosome (Finnegan et al., 2019). These phenotypes highlight the breadth of meiotic drive-induced costs, emphasising the need to investigate the underlying metabolic consequences of harbouring drive across both sexes.

Three diverse measures of metabolic function were assessed to examine the impact of meiotic drive on metabolic life history. Mitochondrial function was measured by monitoring oxygen consumption in thoracic tissue using an O2k-Fluorespirometer (Gnaiger, 2020; Simard et al., 2018). Whole-organism resting metabolic rate was assayed through CO_2_ production measurements using a MAVEn-FT (Multiple Animal Versatile Energetics Flow-Through System, Sable Systems International, Las Vegas, NV). Finally, nutrient acquisition under resting conditions was assessed through the consumption of different diets using a CApillary FEeder (CAFE) assay (Ja et al., 2007). The primary aim was to compare flies carrying drive and wild-type chromosomes, as well as between heterozygous and homozygous females, to test the prediction of general deleterious effects associated with the drive X chromosome. The hypothesis that dysfunction caused by meiotic drive during spermatogenesis leads to direct metabolic consequences was tested by comparing the sexes, with the expectation that males would exhibit more pronounced effects. The hypothesis that there is an accumulation of sexually antagonistic effects linked to drive was tested in the same way but with the reverse expectation that female metabolism would be more strongly impacted.

## 2. Methods and Materials

### 2.1 Stock Generation and Maintenance

The wild-type, standard (ST) stock was collected in 2005 from the Ulu Gombak Valley, Peninsular Malaysia (Cotton et al., 2014; Cotton et al., 2010). Flies with the X^SR^ genotype were collected in 2012 from the same location (Cotton et al., 2014) and, since 2019, have been maintained as a homozygous SR stock as set out in (Finnegan et al., 2019). Experimental ST males (X^ST^/Y) and ST females (X^ST^/X^ST^) were collected on egglays (petri dishes with a damp cotton pad and pureed sweetcorn as food) from cages housing X^ST^/X^ST^ females and X^ST^/Y males. The egglays were incubated at 25°C, and the emerging flies were collected. The same procedure was followed to collect SR males (X^SR^/Y) and heterozygous females (X^SR^/X^ST^) from cages housing X^SR^/X^SR^ females and X^ST^/Y males and to collect homozygous females (X^SR^/X^SR^) from cages housing X^SR^/X^SR^ females and X^SR^/Y males.

### 2.2 Measuring mitochondrial function through high-resolution respirometry

The preparation of *T.dalmanni* thoracic tissue for mitochondrial function analysis was adapted from published methods (Simard et al., 2018; Teulier et al., 2016). Whole thoraces were dissected in 1.5 mL ice-cold MiR05 respirometry buffer (0.5 mM EGTA, 3 mM MgCl2.6H2O, 60 mM lactobionic Acid, 20 mM taurine, 10 mM KH2PO4, 20 mM HEPES, 110 mM D-sucrose, 1 g/L BSA, pH 7.1). The muscle fibres were lightly homogenised in 150 μL of MiR05, of which 50 μL were added in a calibrated O2k-Fluorespirometer (Oroboros Instruments, Innsbruck, Austria) with 2 mL MiR05. The remaining homogenate was frozen and kept for subsequent protein content analysis using a QuantiPro BCA Assay Kit (Sigma-Aldrich).

Pyruvate (10mM) and malate (2mM) (Complex I substrates) were then added to the Oroboros sample chamber, and the LEAK state was recorded after 15–20 minutes. ADP (5mM) was added to reach the OXPHOS state. Cytochrome *c* (10mM) was added to assess mitochondrial membrane integrity, and samples with >20% increased O_2_ consumption were discarded. Glutamate (10mM), succinate (10mM), and glycerophosphate (10mM) were added sequentially, recording O_2_ fluxes at each state. Maximum uncoupled respiration was determined using 0.5μM FFCP. Respiration was inhibited by adding rotenone (0.5 μM), followed by malonic acid (5mM), and antimycin A (2.5 μM), estimating the residual oxygen consumption (ROX). ROX was set as the baseline (O2 consumption = 0), and other states were compared to ROX-corrected states. Finally, Complex IV activity was measured using ascorbate (2 mM) and TMPD (0.5 mM), followed by inhibition with sodium azide (100 mM).

Following Oroboros runs, both respiratory control ratio (RCR) and Complex I-linked efficiency were calculated. The RCR was calculated by dividing the oxygen consumption when ADP was added by the LEAK state (State 3 / State 4). A high RCR value indicates a high mitochondrial coupling of electron transfer to O_2_ reduction and a high respiratory capacity available to phosphorylate ADP to ATP (Gnaiger, 2020).The activity of respiratory Complex I was calculated as the percentage decrease upon the addition of the inhibitor rotenone in the uncoupled state.

Each Oroboros instrument is equipped with two chambers, and two instruments were available for this experiment, enabling the simultaneous testing of four genotypes. Due to this logistical limitation, only wild-type and drive males, as well as wild-type and homozygous drive females, were included in this study. Heterozygous females were not tested in this experiment.

### 2.3 Measuring whole-organism metabolic rate

Whole fly resting metabolic rate was measured using a MAVEn-FT (Multiple Animal Versatile Energetics Flow-Through System; Sable Systems International, USA). Individual flies (average age approximately 50 days post-eclosion) were weighed and then placed into metabolic chambers coupled to an external CO_2_ analyser (LICOR 850; LI-COR, USA). Fly activity was measured via the presence of 3 infrared beams below each chamber, recording movement every time the beams were broken. The chambers (length: 3cm, diameter: 1.3cm) were large enough so that flies could move around but small enough that they could not undertake to exert behaviours such as flying or foraging. The CO_2_ concentration in each chamber was measured during airflow for 120 seconds at a flow rate of 30 mL/min. The system monitored 16 individual chambers consecutively, with three or four measurements for each chamber, over a 3-hour cycle.

The multiplexed configuration included 15 chambers containing flies and one chamber left empty as a control to confirm that there was no interference from extraneous variables. Eight separate trials were conducted, each involving a different set of 15 individuals. Three individuals of the five genotypes were tested in each trial: two male genotypes, X^ST^Y and X^SR^Y, and three female genotypes, X^ST^X^ST^, X^SR^X^ST^ and X^SR^X^SR^. At the beginning of a trial, all individuals were lightly anaesthetised with 5 ppm of CO_2_, weighed (mg), and then allowed to wake and acclimate for 30 minutes before being placed inside the MAVEn-FT system.

### 2.4 Measuring dietary consumption

Food consumption was measured using a CApillary FEeder (CAFE) assay (Ja et al., 2007) adapted for use in stalk-eyed flies. Test individuals were placed individually in vials (20 mL) containing a base layer (6 mL) of 0.8% agar to provide a source of hydration with no caloric content. The top of the tube was secured with a sponge bung penetrated by a pipette tip cut to fit a capillary tube. Flies remained in their agar vials overnight. The following morning, a 20 μL glass capillary filled with the allocated dietary treatment was inserted into each individual vial (SI Figure 1). Dietary treatments used in this experiment were a mix of microbiology yeast extract and sugar, but all with a final concentration of 32.5 g/L. The three diet treatments used were high protein (1:1 yeast:sugar), low protein (1:2 yeast:sugar) and sugar alone (10% solution). Blank controls (empty vials containing a liquid capillary but with no fly) were used as controls to estimate the volume of liquid lost through evaporation alone.

**Figure 1:**
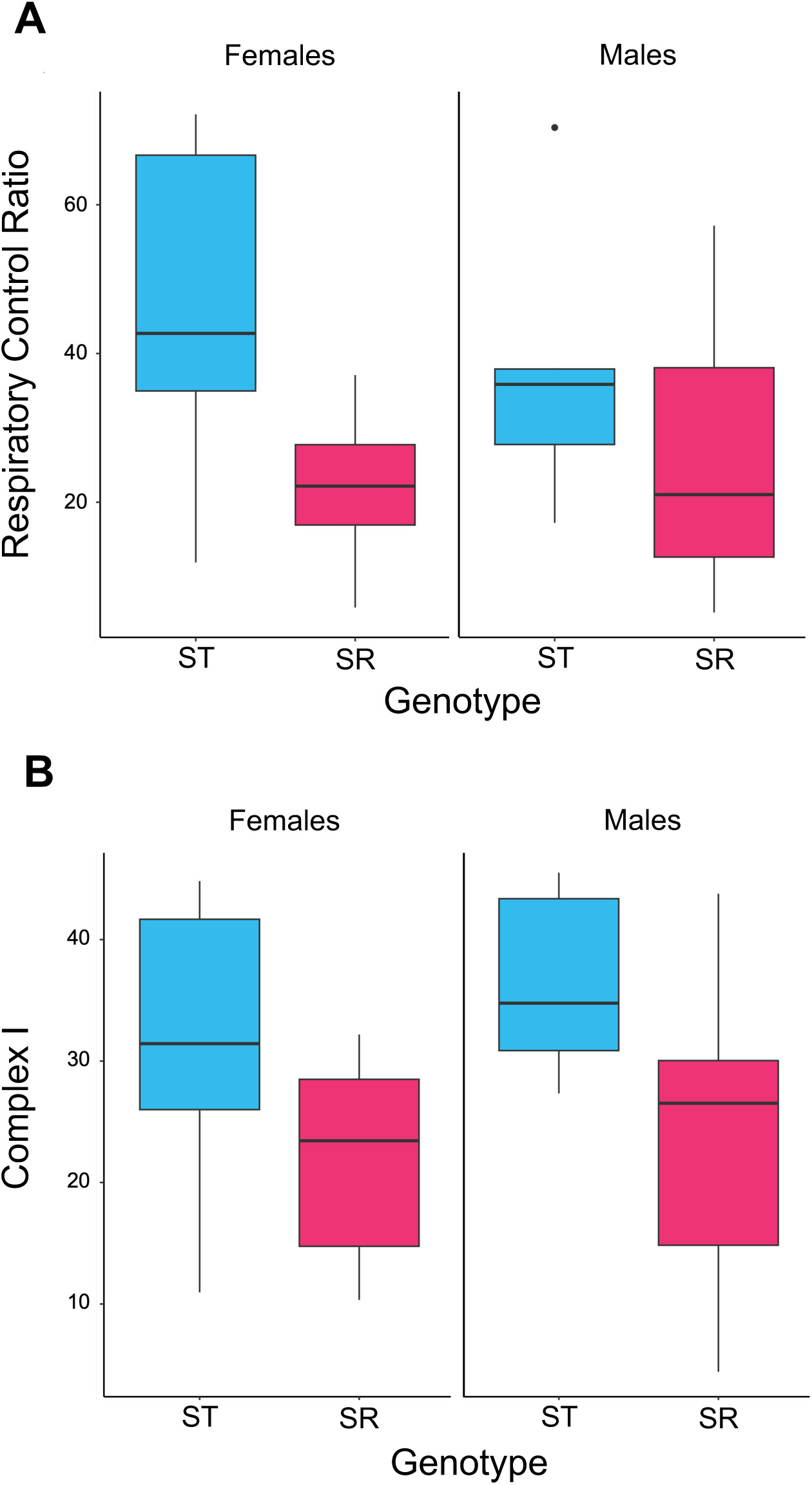
A) Respiratory Control Ratio and B) percentage of Complex I contribution to respiration in females (left) and males (right), for ST (blue) and SR (pink) genotypes. Females with SR genotype were SR homozygotes. The central line in each box represents the median, the box indicates the interquartile range and the whiskers are the 95% confidence interval.

All vials were placed in an incubator (PHCI - MLR-352H Climate Chamber) at 25 °C and 90% humidity to maintain moisture in the vials and minimise capillary liquid evaporation. The liquid meniscus at the top of the capillaries was marked, and the decline over known time periods due to consumption and evaporation was recorded. The physical distance between these marks was measured using digital callipers to obtain the volume of liquid consumed (μL) from the 20 μL capillary. Consumption was measured, and food was replaced after 20h, 40h and 60h, resulting in the total consumption (calculated as the sum across the three days).

The five genotypes were tested, two male and three female, as listed in section 2.3. The thorax of ice-anaesthetised flies was measured upon completion of the experiment using an Infinity Capture video microscope attached to a computer equipped with NIH image software (FIJI - IMAGEJ, version 2.1.0/1.53c). The thorax was measured from the prothorax anterior tip along the midline to the joint in between the thorax and metathoracic legs (Rogers et al., 2008).

### 2.5 Statistical Analyses

All statistical analyses were carried out using R Studio (Version 2023.12.0). We used linear models using either the “*lm*” or “*lmer*” function in R, depending on whether the models had random effects.

#### Mitochondrial function

Oxygen consumption data was normalised by protein content within the Oroboros software (DatLab v7.8). Sex and genotype were accounted for in all models as fixed effects. Interactions between variables were tested, and the model was reduced by removing interaction terms that were not significant. Response variables were RCR and Complex I function.

#### Whole-organism metabolic rate

MAVEn data files were processed using Sable Systems software (ExpeData v.1.7), which extracted the 30-second window (within the 2-minute reading) with the lowest CO_2_ production and corresponding activity values of individual flies. Respiration was corrected by weight in all models to obtain the mass-independent metabolic rate. Sex, activity level, age of fly and genotype were accounted for in all linear regression models. Interactions between variables were tested, and the model was reduced by removing interaction terms that were not significant. Given we obtained three to four measurements from the same individual, we added the chamber replicate measurement nested within each individual as a random effect.

Genotype was assessed using two methods. In the first, the five genotypes were reduced to two categories, ST or SR, for each sex. The SR category joined together females with at least one drive chromosome, that is, heterozygotes (X_SR_/X_ST_) and drive homozygotes (X_SR_/X_SR_), allowing an easier comparison of females and hemizygous males. Alternatively, the full set of female genotypes was considered to consider whether there was evidence of dominance.

#### Dietary consumption using the CAFE assay

Prior to the analysis of the data, we corrected total liquid food consumption by mean evaporation rate. This involved subtracting the mean food loss due to evaporation (blank vials with no fly) from all experimental vials with flies. The data were log-transformed to meet statistical model assumptions of normality. The full models included consumption values as a response variable, with thorax (as a proxy for body size), sex, treatment (diet type) and genotype as fixed factors. Interactions between variables were tested, and the model was reduced by removing interaction terms that were not significant. The random effect of “box” was also included to control for any batch differences across experimental runs. Genotype was assessed using the two methods in the whole organism metabolic rate subsection.

### 2.6 Replication Statement

## 3. Results

### 3.1 Mitochondrial function is genotype-dependent

Genotype affected mitochondrial activity, as measured by the respiratory control ratio (*F*_2,23_ = 6.394, *p* = 0.019), where ST individuals had higher RCR (42.540 ± 5.669) than SR individuals (24.375 ± 4.201; Figure 1A). There was no difference across the sexes (*F*_2,23_ = 0.151, *p* = 0.701) and no interaction between genotype and sex (*F*_3, 22_ = 0.995, *p* = 0.329).

In addition, ST individuals displayed a higher Complex I contribution to respiration (33.902 ± 2.815) than SR individuals (22.762 ± 3.180; *F*_2,23_ = 6.741, *p* = 0.016; Figure 1B). Again, there was no difference across the sexes (*F*_2,23_ = 0.512, *p* = 0.482) and no interaction between genotype and sex (*F*_3,22_ = 0.115, *p* = 0.738). All other respiratory states of the electron transport system did not vary between SR and ST individuals (*p* > 0.05, SI Table 1).

**Table 1:** Table comprising replication units for all experiments.

| **Experiment** | Scale of inference | Scale at which the factor of interest is applied | Number of replicates at the appropriate scale |
| --- | --- | --- | --- |
| Mitochondrial function | Genotype, per sex | Individual stalk-eyed flies | 6–7 |
| Whole-organism metabolic rate | Genotype, per sex | Individual stalk-eyed flies | 24–36 |
| Whole-organism metabolic rate | Females, per genotype | Individual stalk-eyed flies | 24 |
| Dietary consumption | Genotype, per sex, per diet | Individual stalk-eyed flies | 12–33 |
| Dietary consumption | Females, per genotype, per diet | Individual stalk-eyed flies | 15–18 |

### 3.2 Whole-organism metabolic rate is sex- and genotype-dependent

Pooling SR heterozygotes and homozygotes, genotype altered respiration (*F*_1,101.2_ = 7.152, p = 0.009), with a higher level of CO_2_ production in individuals carrying SR chromosomes (1.846×10^−4^ ± 4.547×10^−6^) compared to those with only ST chromosomes (1.895×10^−4^ ± 6.025×10^−6^). Respiration also differed between the sexes (*F*_1,123.2_ = 5.110, *p* = 0.026), with males (1.958×10^−6^ ± 6.100×10^−6^) having greater respiration than females (1.803×10^−6^ ± 4.471×10^−6^). There was no interaction between genotype and sex (*F*_1,124.6_ = 1.042, *p* = 0.309).

Limiting the analysis to females, respiration differed amongst the three genotypes (*F*_2,78.6_ = 6.241, *p* = 0.003, Figure 2). SR homozygous females showed the same level of respiration as heterozygotes (Tukey’s HSD: *t* = −0.032, *p* = 0.999, df = 66.8; Figure 2). Both SR homozygotes (*t* = 3.106, *p* = 0.008, *df* = 73.300) and heterozygotes (*t* = 3.034, *p* = 0.010, *df* = 56.900) had elevated respiration compared to ST homozygotes. In addition, there was no difference in the cross-sex comparison of all individuals with drive chromosomes (SR males, female heterozygotes and female SR homozygotes; all *t* < 1.182, *p* > 0.468).

**Figure 2:**
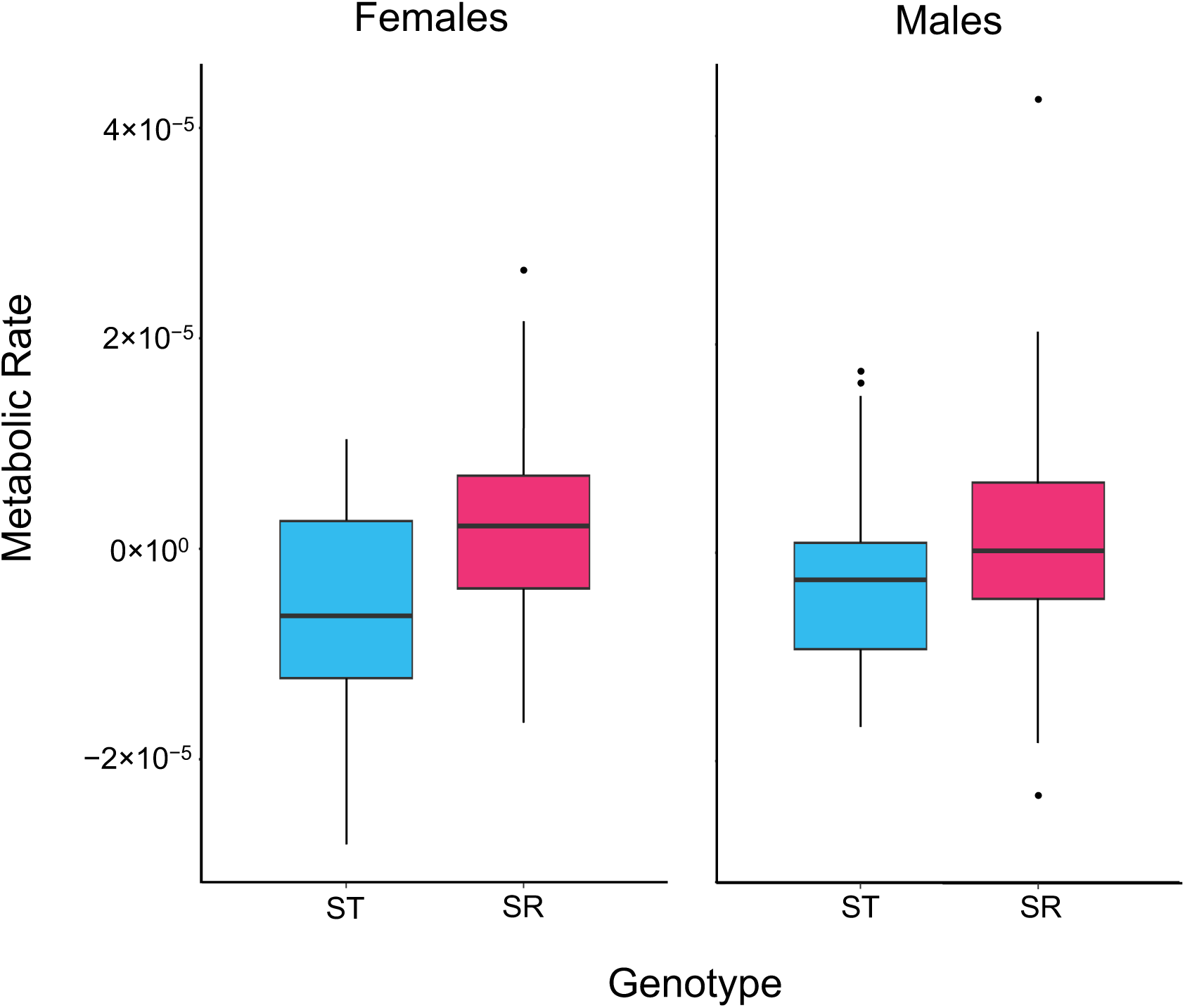
CO_2_ produced as a measure of metabolic rate in females (left) and males (right), for ST individuals (blue) and SR individuals (pink). Residual metabolic rate is plotted after accounting for body size variation. Heterozygotes and SR homozygous females are pooled. The central line in each box represents the median, the box indicates the interquartile range and the whiskers are the 95% confidence interval.

### 3.3 Dietary consumption using the CAFE assay is diet- and genotype-dependent

Pooling SR heterozygotes and homozygotes, genotype altered food consumption (*F*_1,224.96_ = 7.966, *p* = 0.005), as individuals carrying SR chromosomes (7.559 ± 0.513) consumed a greater total amount of food than individuals with only ST chromosomes (4.855 ± 0.520). There was no effect of sex on food consumption (*F*_1,223.68_ = 3.377, *p* = 0.067). Consumption varied with diet (*F*_2,224.01_ = 21.401, *p* < 0.001), with more of the high protein diet consumed than the low protein diet (*t* = 2.747, *p* = 0.018, *df* = 225), which in turn was consumed more than the sugar diet (*t =* 3.784, *p* < 0.001, *df* = 224; Figure 3). There was no interaction between diet and genotype (*F*_2,222_ = 0.264, *p* = 0.768) or between diet and sex (*F*_1,223.53_ = 0.033, *p* = 0.855).

**Figure 3:**
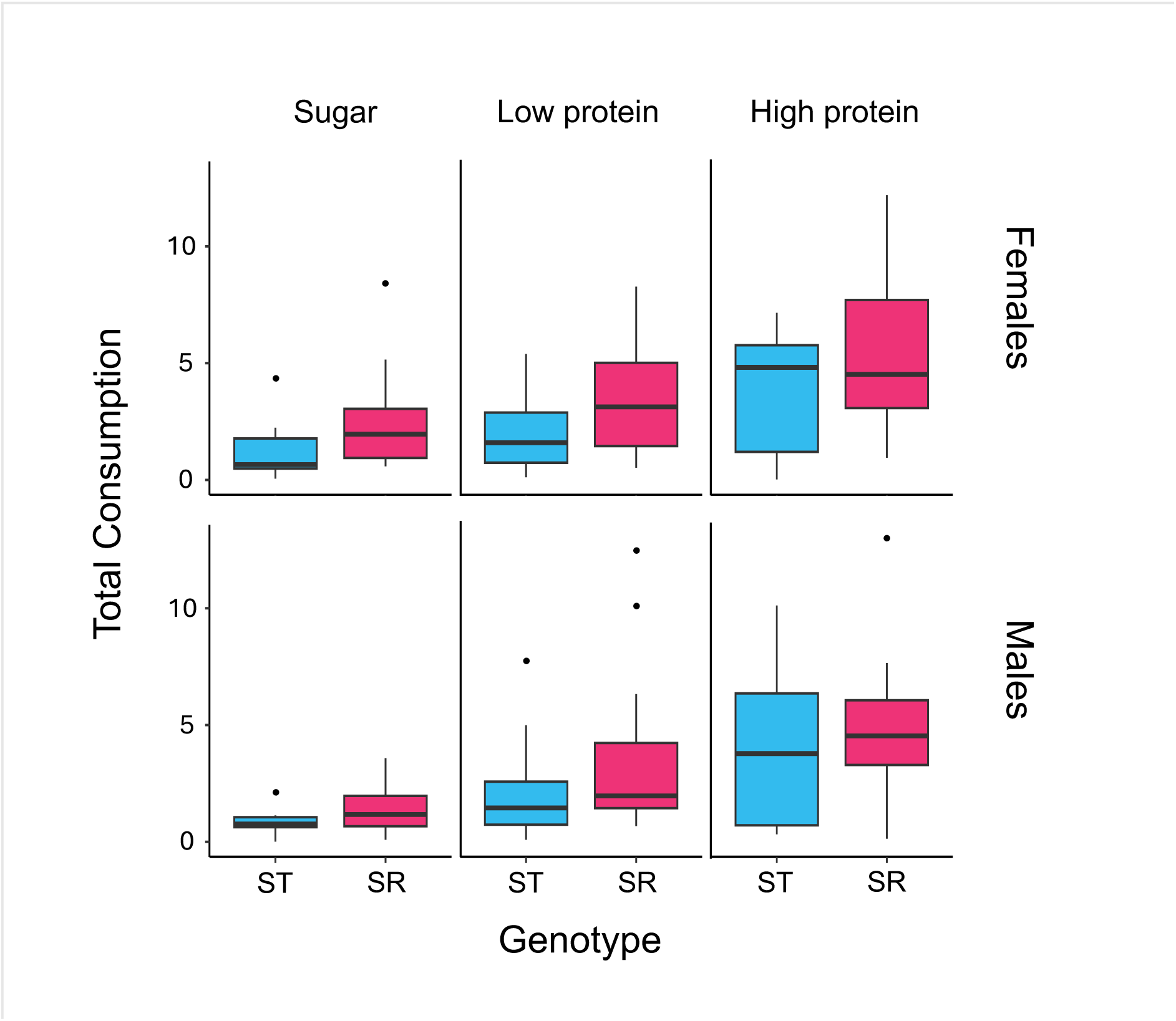
Total consumption (corrected for body size) of sugar (left), low protein (middle) and high protein (right) by females (top) and males (bottom). Genotypes are ST (blue) and SR (pink). Heterozygotes and SR homozygous females are pooled. The central line in each box represents the median, the box indicates the interquartile range and the whiskers are the 95% confidence interval.

Limiting the analysis to females, consumption differed amongst the three genotypes (*F*_2,137.22_ = 3.726, *p* = 0.027). Post-hoc comparisons showed that heterozygotes consumed more than ST homozygotes (*t* = −2.648, *p* = 0.024, df = 138), but other comparisons were not significant. Consumption again varied with diet (*F*_2,136.97_ = 10.900, *p* < 0.001). The high protein diet was consumed more than the sugar diet (*t* = 4.665, *p* < 0.001, *df* = 137). But there was no difference between the consumption of high protein and low protein diets (*t =* 2.359, *p =* 0.051, *df* = 137) or low protein and sugar diets (*t =* 2.324, *p =* 0.056, *df* = 137), though both comparisons were borderline significant. There, again, was no interaction between diet and genotype (*F*_4,132.99_ = 0.267, *p* = 0.899). In addition, there was no difference in consumption for the cross-sex comparison of all individuals with drive chromosomes (SR males, female heterozygotes and female SR homozygotes (all *t* < 1.643, *p* > 0.231).

## 4. Discussion

The X-linked sex ratio (SR) meiotic drive system in stalk-eyed flies has distinct metabolic consequences. We provide evidence that drive flies exhibit mitochondrial dysfunction, and this dysfunction is compensated via an increase in basal respiration rate. They also consume more food per unit of time across a range of diets, indicating further compensatory mechanisms to counteract their less efficient metabolism and a potential inability to sufficiently utilise nutrients. These results provide the first demonstration of how metabolic function can be corrupted by meiotic drive. All the markers of stress within the metabolic system occurred in females as well as males carrying the drive chromosome, despite the direct disruption of meiosis in stalk-eyed flies being limited to spermatogenesis. This strongly points to the genomic inversions on the driving X chromosome being linked to a raised mutational load, which is causing the range of metabolic costs measured.

Mitochondrial function was measured through high-resolution respirometry using an O2k-Fluorespirometer (Oroboros Instruments, Austria). This measures the flux through the electron transport chain during OXPHOS using a substrate-uncoupler-inhibitor titration protocol (see SI for details). Two aspects were found to be compromised. Individuals carrying meiotic drive had a lower respiratory control ratio (RCR) (Figure 1A), indicating the uncoupling of oxidative phosphorylation from ATP synthesis (Divakaruni et al., 2022). They also presented reduced Complex I substrate oxidation contribution to respiration (Figure 1B). Complex I is the first and largest unit in the respiratory chain and the primary contributor to the proton motive force that facilitates ATP synthesis (Sharma et al., 2009). Other mitochondrial states tested were not found to differ between SR and ST individuals (SI Table 1). A possible cause of low RCR and Complex I dysfunction is mutations in the nuclear DNA encoding mitochondrial subunits or accessory proteins located within the driving X chromosome inversions. For example, several X-linked mutations associated with Complex I dysfunction have been characterised by various pathologies (Fassone et al., 2012; Reinson et al., 2019), including the NDUFA1 gene implicated in mitochondrial encephalomyopathy (Fernandez-Moreira et al., 2007). These possibilities could be tested in future bioinformatic work comparing SR and wild-type sequence data (Reinhardt et al., 2023), specifically looking for evidence of disruption to coding sequences of nuclear genes involved in mitochondrial function or for differential expression of nuclear and mitochondrial genes involved.

There were no sex differences in the RCR or Complex I measurements, but the sample size was not large, which limits the assurance of this result. However, it is noteworthy to mention that the number of samples tested in this experiment is typical of protocols used in this field (Gonçalves et al., 2022; Nääv et al., 2020; Pichaud et al., 2013; Rodríguez et al., 2021, 2023). A potential shortcoming is that the O2k assessment was limited to female homozygotes, meaning that the degree of dominance of drive effects on mitochondrial activity in females cannot be discerned. This could be investigated as part of future work. Another area to look at is male reproductive tissues where the drive genes causing disruption of spermatogenesis are active (i.e. the testes and accessory glands (Reinhardt et al., 2014)). These tissues are expected to show elevated dysfunction compared to thorax musculature due to the direct effect of drive, which might be measurable through mitochondrial deficits in RCR, Complex I or other elements of the respiratory chain. Previous studies in the *t* haplotype system in mice provide valuable parallels. Mutant *t^n^* sperm show increased aerobic metabolism (exemplified by a reduced NADH/NAD ratio), which is thought to put them at a selective advantage because it increases motility, maturation and fertilisation (Ginsberg et al., 1974). Our study used thoracic tissues, but future experiments on reproductive tissues would provide further insight. In some instances of sperm-killer drive, some drive sperm experience pleiotropic collateral damage (Price et al., 2008). However, it was recently found that SR *T. dalmanni* males show no reduction or advantage in sperm competitiveness or paternity (Bates, 2023; Bates et al., 2023). The most plausible explanation is the compensatory enlargement of SR male testes, with a concomitant equal sperm count per ejaculate when compared to wild-type males (Meade et al., 2020; Meade et al., 2019). A further possibility that remains to be investigated is whether there are alterations to the metabolic rate of SR males. In contrast, female reproductive tissues are not expected to be differentially targeted compared with somatic tissues, as meiotic drive has no known downstream consequences for oogenesis in stalk-eyed flies.

The whole-organism metabolic rate of stalk-eyed flies was examined using a MAVEn-FT system, which takes repeated measurements of CO_2_ production of resting flies over a 3-hour period. We found whole organism respiration to be higher in males than females and also higher in drive individuals within each sex. These findings suggest that less efficient mitochondrial function induces a compensatory elevated metabolic rate in drive individuals. Furthermore, drive heterozygotes and homozygotes were tested alongside wild-type homozygous females for metabolic rate, which demonstrated that the SR chromosome has a dominant effect on respiration, as females with a single SR chromosome produced as much CO_2_ as those with two (Figure 2). Likewise, there was no difference between drive males and the two female genotypes carrying drive (Figure 2). These results are consistent with the notion that most of the metabolic costs arising are associated with the mutational load in the SR chromosome inversions, as was seen in direct measurements of mitochondrial activity. Similar increases in CO_2_ production have been noted in other systems, like *Drosophila melanogaster* when there is an incompatibility between the nucleus and the mitochondria (Bettinazzi et al., 2024; Hoekstra et al., 2013). However, opposing results were found in mice, as females carrying the *t* locus meiotic drive system were found to exhibit a lower resting metabolic rate compared to wild-type mice in larger individuals (Lopes et al., 2019). In mice, a lower metabolic rate has been linked to increased longevity (Duarte et al., 2014). This association leads the authors to argue that reduced metabolism is an adaptation in drive females that compensates for the smaller litter sizes seen in *t* drive females (Sutter et al., 2015) by extending the number of litters that they sire during their lifespan (Duarte et al., 2014). The decrease in metabolic rate with body size is not observed in males carrying the *t* locus, so it is not a direct consequence of drive; rather, it is a female-only phenomenon, which aligns with the adaptation hypothesis (Duarte et al., 2014).

A final test of metabolic costs arising through meiotic drive measured food intake over a range of diets using the CAFE assay. Food consumption is a good proxy for metabolic-reliant phenotypes, given its involvement in fitness and lifespan across many species, including *D. melanogaster* (Camus et al., 2019; Skorupa et al., 2008), and its association with metabolic rate (Nicholls et al., 2021; Roark et al., 2009; Winwood-Smith et al., 2017). SR individuals consistently consumed more, with no difference between females and males (Figure 3). Consumption varied markedly with diet, and intake increased with greater amounts of protein in the diet provided. However, diet type had no impact on the difference between SR and wild-type flies. This suggests that metabolic deficiency amongst flies carrying drive chromosomes is compensated by greater food intake. Even though the value of different food sources varies with the reproductive demands of females and males (Camus et al., 2018; Reddiex et al., 2013), this was not observed in this assay. Typically, females prefer diets with greater protein content (for egg production), and, in contrast, males prefer carbohydrates to power male-male competition and sexual display (Jensen et al., 2015; Kodric-Brown et al., 1987; Maklakov et al., 2008; Wheeler, 1996). Even if this is the case in stalk-eyed flies, it did not follow through to cause differentiation of consumption in flies with drive chromosomes or across the sexes. Drive individuals did not show consumption modification in response to their mitochondrial dysfunction across different diets other than an overall increase in consumption. In addition, as before in the whole-organism respiration tests, there was no difference between female heterozygotes and homozygotes, indicating dominance of the dysfunction caused by meiotic drive. Nor did these two female genotypes differ in consumption compared to drive males (Figure 3).

Altogether, these results point to the preponderance of indirect mutational load costs being linked to meiotic drive in a sex-independent manner. This does not rule out the existence of direct effects of meiotic drive in males that reduce metabolic function, nor the possibility of selection for linked sexually antagonistic alleles that depress female metabolic function (Rydzewski et al., 2016). What it suggests is that these effects are relatively less important and subsumed by the indirect mutational load held in the multiple inversions that cover almost all of the SR X chromosome in *T. dalmanni* and other examples of X-linked drive (Jaenike, 2001; Paczolt et al., 2017). Metabolic function measured in this study concerned somatic tissue, whole-organism respiration and dietary consumption. It may be that reproductive tissues where meiotic drive takes place (i.e. male testes) or gametogenesis in general (i.e. including ovarian tissue) have different metabolic responses reflecting the direct effect of meiotic drive genes or sexually antagonistic gene expression, respectively. This possibility suggests future work.

## 5. Conclusions

This research provides the first evidence of the range of metabolic costs associated with a meiotic drive system in stalk-eyed flies. The costs uncovered include mitochondrial dysfunction (lowered respiratory control ratio and reduced Complex I contributions to respiration), metabolic inefficiency (higher basal metabolic rate) and increased food consumption across a range of diets (varying in protein content). The latter two are likely to be compensatory mechanisms arising from mitochondrial dysfunction. The costs were of equal scale across the sexes even though the direct action of meiotic drive only manifests as a loss of gametes in male spermatogenesis. This supports the hypothesis of predominant indirect costs associated with meiotic drive. These are likely to arise from genes linked to meiotic drive on the X chromosome. The driving X chromosome contains a number of large genomic inversions, which restrict its recombination with the wild-type X chromosome. As the meiotic drive X chromosome is at low frequency in natural populations (∼20%), it is subjected to weak selection and the accumulation of deleterious mutations which impact metabolic function. By comparing heterozygous and homozygous females with hemizygous males, the analysis found no evidence of predominant recessive effects in females as predicted by sexually antagonistic selection. Surprisingly, there was little evidence of greater metabolic deficits in males. While many life history phenotypes have been examined in the *T. dalmanni* system, future work will aim to strengthen what is known about metabolic phenotypes that potentially give rise to these traits. A further analysis of male reproductive tissues is needed to evaluate whether male-specific metabolic costs are evident where the meiotic drive genes themselves are active. Taken together, these results offer new insights into the metabolic and energetic underpinnings of harbouring meiotic drive systems.

## Acknowledgements

SLB is funded by a Studentship from the London NERC DTP (NE/S007229/1), POM is supported by funding from the Engineering and Physical Sciences Research Council (EP/F500351/1, EP/I017909/1), Natural Environment Research Council (NE/R010579/1, NE/X009734/1) and Biotechnology and Biological Sciences Research Council (BB/V003542/1). MFC is funded by a Natural Environment Research Council Fellowship (NE/V014307/1) and a Leverhulme Trust grant (RPG-2023-198). CTY and CDV were supported by the Harold & Olga Fund Summer Studentships. We thank Rebecca Finlay and Wendy Hart for helping with the maintenance of stalk-eyed fly stocks, and Nick Lane for discussion.

## Author Contributions

SLB, POM & MFC conceived the ideas and designed the methodology; SB, ER, & HW collected the mitochondrial data, SB, CTY, CDV & MFC collected the respiration data, SB & AH collected the dietary consumption data; SLB, ER, POM & MFC analysed the data; SB, POM & MFC led the writing of the manuscript. All authors contributed critically to the drafts and gave final approval for publication.

## Extended Methods

### Mitochondrial function measurement through high-resolution respirometry

#### Sample preparation

The preparation of *T. dalmanni* thoracic tissue for mitochondrial function was developed following previously published methods (Teulier *et al*., 2016; Simard *et al*., 2018). Whole thoraces from individual *T. dalmanni* were weighed and transferred to a 24-well plate filled with 1.5 mL of ice-cold MiR05 respiration buffer (0.5mm EGTA, 3mm MgCl_2_·6H_2_O, 60 mM lactobionic acid, 20mm taurine, 10mM KH_2_PO_4_, 20mm HEPES, 110mm D-sucrose and pH 7.1) for dissection (Bettinazzi *et al*., 2019). The cuticle was removed, and the tissue was transferred into a 1.5 mL Eppendorf tube containing 150 μL MiR05 for homogenisation. Gentle manual homogenisation was performed using a polypropylene Kimble Kontes Pellet Pestle, and 50 μL of the tissue homogenate was transferred to a 2 mL respiration chamber of a high-resolution O2k fluorespirometer (Oroboros Instruments, Innsbruck, Austria), previously calibrated at 25°C with 2 mL of the respiratory medium MiR05.

#### Initial respiration assessment

After adding the tissue homogenate, Complex I substrates pyruvate (10 mM) and malate (2 mM) were injected, and the chambers closed. After 15–20 minutes of signal stabilisation, the respiration of the LEAK (N_L) state was recorded. ADP (5 mM) was then added to reach the OXPHOS respiratory state sustained by Complex I (N_P) to determine the OXPHOS capacity.

#### Checking sample integrity

Cytochrome *c* (10mM) was added to check the sample preparation quality, which reflects the integrity of the mitochondrial outer membrane and was thus used to assess the quality of mitochondrial sample preparation. Samples where O_2_ consumption of the prepared homogenate increased by more than 20% after cyt *c* injection were discarded.

#### Sequential substrate addition and respiration measurement

Next, glutamate (10 mM, NG_P state), succinate (10 mM, NGS_P state), and glycerophosphate (10 mM, NGSGp_P state) were added sequentially, and O_2_ fluxes were recorded at each stable state, with the last addition yielding the maximum coupled respiration. Maximum uncoupled respiration (NGSGp_E) was then evaluated using 0.5 μM of the uncoupler FCCP (1 µl step titration) until it attained a plateau indicating maximal ETS capacity.

#### Sequential inhibition

Respiration was then inhibited by stepwise additions of the Complex I inhibitor rotenone (0.5 μM, SGp_E), the Complex II inhibitor malonic acid (5 mM, Gp_E) and the Complex III inhibitor antimycin A (2.5 μM). The residual oxygen consumption (ROX) could then be estimated to correspond to the O_2_ consumption due to oxidative side reactions after inhibiting the ETS (Gnaiger, 2020). ROX was corrected as the baseline state (i.e., O_2_ consumption = 0 at ROX), and all other respiratory states (e.g., N_L, N_P, NGSGp_P, etc.) were compared to the corresponding ROX-corrected state for statistical analysis.

#### Reoxygenation and Complex IV assessment

Reoxygenation before Complex IV assessment is necessary as its substrates rapidly consume oxygen. The chambers were opened for reoxygenation and then closed to measure Complex IV activity using ascorbate (2 mM) and TMPD (0.5 mM). Finally, the Complex IV was inhibited by injecting 100 mM sodium azide.

#### Protein quantification

Results are expressed as the O_2_ consumption per protein content (mg/mL) of the thorax homogenate. Protein assays were performed on each individual’s leftover homogenate using a QuantiPro^TM^ BCA assay kit (Merck KGaA, Darmstadt, Germany) following the manufacturer’s instructions. Several parameters were calculated to show the functioning of mitochondria during respiration.

#### Data processing

Raw data was extracted from DatLab (version 7.4), processed and analysed using calculation templates provided by the manufacturer. The respiratory acceptor control ratio (RCR) was calculated as RCR = N_P/N_L to show the efficiency of OXPHOS coupling during respiration. Substrate contributions to the respiration were represented as the percentage increase in O_2_ consumption upon the addition of each substrate (glutamate: NG_P – N_P/N_P*100; succinate: NGS_P – NG_P/NG_P*100; G3p: NGSGp_P – NGS_P/NGS_P*100). The activity of each respiratory Complex was calculated as the percentage decrease upon the addition of each inhibitor in the uncoupled state (e.g., CI: NGSGp_E – SGp_E/NGSGp_E*100 and CII: SGp_E – Gp_E/SGp_E*100). Moreover, flux control ratios (FCR) were calculated using the maximal ETS capacity as the reference state (NGSGp_E) to allow internal comparison to respiratory states and limitation of OXPHOS capacity by the phosphorylation system.

